# Relatedness and last male sperm precedence in *Cheilomenes sexmaculata*: Implications for inbreeding avoidance

**DOI:** 10.64898/2026.10.15.759354

**Authors:** Swati Saxena, Shriza Rai, Geetanjali Mishra, Omkar

**Affiliations:** Research Scholar, University of Lucknow, India

**Keywords:** Inbreeding avoidance, ladybird, mate discrimination, sexual selection, mate preference, paternity share, siblings, reproductive strategies

## Abstract

Avoidance to mate with relatives is a common evolutionary strategy, as it helps maintain genetic diversity and overall fitness within populations. Females often prefer unrelated males to mitigate the risks associated with inbreeding, such as the expression of deleterious recessive alleles. This preference can involve various post-copulatory mechanisms through which females bias paternity towards males with potentially better genes. The phenomenon of last male sperm precedence, where the last male to mate with a female tends to sire the majority of offspring, is well-documented in ladybird beetles. However, how relatedness affects this phenomenon remains unclear. This study aims to investigate whether paternity share is altered when the last male to mate with a female is related to her. By conducting double mating experiments with two different morphs (typical and intermediate) as phenotypic markers, we seek to determine if relatedness disrupts last male sperm precedence. It was hypothesised that females of *Cheilomenes sexmaculata* will prefer unrelated males over related ones, irrespective of mating order, potentially breaking down the last male sperm precedence. This preference for unrelated males was reflected in our results with unrelated males exhibiting enhanced reproductive success. Females were more fecund and exhibited higher egg viability with unrelated males. Greater paternity share was observed for unrelated males when the second male was related to the female. This study highlights the dynamic nature of sperm precedence and the role of mate quality in female mate choice, emphasizing the importance of inbreeding avoidance in sexual selection.

## Introduction

Parker’s (1970) landmark paper on sperm competition expanded sexual selection beyond mating. Post-copulatory sexual selection, which affects both mating performance and ejaculate quality, is widespread due to the prevalence of polyandry (Taylor et al., 2014; Thornhill and Alcock, 1983). The good genes model suggests individuals with superior genetic quality achieve greater reproductive success. Mechanisms like mate selection and competitive interactions favour those with better genetic attributes (Zahavi, 1975; Hamilton and Zuk, 1982; Houle and Kondrashov, 2002; Kokko et al., 2002, 2003; Fedina and Lewis, 2007; Whitlock and Agrawal, 2009).

Sexual selection favours traits that help individuals choose mates based on their reproductive state, enhancing success (Parker, 1979; Rosenthal and Ryan, 2022). Mating and gamete investment are strategically adjusted due to the costly and limited nature of sperm and ova (Dewsbury, 1982; Arnqvist and Nilsson, 2000; Moore, 2014; Macartney et al., 2021), leading to promiscuous males and choosy females (Trivers, 1972; Andersson, 1994). Few studies have investigated the link between female premating choices and the outcomes of copulatory and post-copulatory processes (Danielsson, 2001; Bussiegre et al., 2007; Demary and Lewis, 2007; Matzke et al., 2023). For instance, *Tribolium castaneum* females use olfactory cues to choose mates and exercise cryptic choice during mating (Fedina and Lewis, 2006; Fedina, 2007). Females of *Poecilia reticulata* prefer colourful males and accept more sperm from them (Evans et al., 2003; Pilastro et al., 2004).

In case of multiple mating, acquisition of higher quality sperm to pass on “better genes” to their offspring remains a challenge. Females choose mates that offer the best fitness benefits, including direct advantages like better foraging territories and indirect benefits from high genetic quality (Halliday, 1983; Berson and Simmons, 2018). They may correct mate choice errors based on cues about male phenotypic or genetic quality during mating (Danielsson, 2001; Demary and Lewis, 2007). While many studies have shown that mating between individuals with similar genotypes increases the homozygosity of deleterious mutations, which can lead to reduced viability—a phenomenon known as inbreeding depression (Charlesworth and Charlesworth, 1987; Keller and Waller, 2002) animals may also select genetically dissimilar mates, leading to fitness benefits, a concept known as optimal outbreeding theory (Bateson 1978, 1983). Since mating between relatives reduces offspring viability and fertility due to inbreeding depression (Charlesworth and Charlesworth 1987; Chen et al., 2017), females avoid inbreeding to secure genetic benefits for their offspring (Jennions and Petrie, 2000; Duthie et al., 2019).

Inbreeding avoidance in insects can occur through pre-copulatory mechanisms, such as avoiding related males in butterflies (Fischer et al., 2015), or through post-copulatory mechanisms, like preferential use of stored sperm from unrelated males in crickets (Bretman et al., 2009). The genetic compatibility hypothesis suggests that females prefer mates with the most compatible genomes to enhance offspring fitness (Trivers 1972; Zeh and Zeh 1996, 1997; Hine et al., 2002). This preference involves various post-copulatory mechanisms that bias paternity towards males with potentially better genes, ensuring higher reproductive fitness for their progeny (Bateson 1983; Simmons 1991; Charlesworth and Willis 2009; Duthie et al., 2019). For example, in *Blattella germanica* (L.), females preferentially mate with non-sibling males over sibling males to avoid inbreeding (Lihoreau et al., 2007).

Ladybird beetles, a potent biocontrol agent (Shandilya et al., 2024), exhibit multiple mating (Haddrill et al., 2008) and are known to display mate choice (Mishra and Omkar 2014; Dubey et al. 2016a, b) and kin selection (Pervez and Khan, 2020). In a study on a species of ladybird beetle, *Cheilomenes sexmaculata* (Fabricius, 1781), Saxena et al (2016) demonstrated that these beetles show bias towards unrelated mates and avoid mating with relatives. Females not only delayed the onset of copula but fecundity and percent egg viability were also found to be reduced following mating with the related males (Saxena et al., 2016). Besides, relatedness also modulated mate preference in *C. sexmaculata* (Saxena et al., 2021). Females showed a preference for polygynous males over monogynous males, suggesting superior fitness in mate search competition, sperm supply, ejaculate quality, and viability. However, this preference reversed when the polygynous males were related to the female, leading them to choose unrelated monogynous males instead. The “good gene” hypothesis suggests that females prefer high-quality or more experienced males for both direct and indirect benefits (Simmons, 2001a). Since, mating between genetically related individuals increases homozygosity, thereby enhancing the risk of inbreeding depression (Müller & Juškauskas, 2018; Müller & Müller, 2016; Valtonen et al., 2014). Hence suggesting that *C. sexmaculata* exhibit inbreeding avoidance to avoid the production of low-quality offspring. However, inbreeding avoidance is not universal (de Boer et al., 2021). Also, through these studies it is not clear how widespread effects of relatedness are on sperm precedence and hence paternity share. As previous studies on ladybird beetle, *C. sexmaculata*, reported to exhibit last male sperm precedence and the last male to mate sires up to 72% of the total offspring produced (Chaudhary et al. 2016), it is not known if the last male sperm precedence is a static or rather a dynamic phenomenon and can be modulated depending upon the quality of mates. Keeping this in view, the present experiment was conducted to test whether relatedness can break down the last male sperm precedence. We hypothesized that *C. sexmaculata* females, being able to distinguish between mates, will show a bias towards unrelated males and prefer them regardless of mating order. This preference is expected to result in greater paternity share for unrelated males when the second male is related to the female. How this preference may disrupt the last male sperm precedence has been accomplished through double mating experiments using different morphs as phenotypic markers. This study will provide insight into the dynamic nature of sperm precedence and the role of mate quality in female mate choice, emphasizing the importance of inbreeding avoidance in sexual selection.

## Materials and Methods

### Experimental organism

*Cheilomenes sexmaculata*, also known as the zig zag ladybird beetle, is a polymorphic (Kawakami et al. 2015; Dubey et al. 2016a), abundantly available, generalist ladybird beetle (Agarwal and Yasuda 2000). These beetles exhibit three morph types based on degree of melanization of elytra: typical, intermediate, and pure melanic (Plate 1). The typical and intermediate morphs are more prevalent in field conditions, while the pure melanic morph is extremely rare and reared strictly under laboratory conditions. Therefore, these beetles can be broadly categorized into two main morphs: typical and melanics, which vary in their degree of melanization (intermediates henceforth). The typical morph has a clear black (melanised) zig zag pattern over the orange-red elytra and the pure melanic morph has a fully melanized elytra. The intermediate morph exhibits a variable degree of melanization such as the zig zag pattern appearing to be fused due to increased melanization along withnone to some melanization on the pronotum. Pairs with the same morph (homomorphic) produce offspring of that same morph. In contrast, pairs with different morphs (hetromorphic), such as typical and melanic, result in offspring that are lighter than the melanic parent but darker than the typical (Dubey et al., 2016) (Plate 2). Therefore, these morphs function as phenotypic markers to ascertain the paternity of the offspring.

### Stock maintenance

Adults of typical (T) and intermediate (I) morphs of *C. sexmaculata* were collected from agriculture fields of Lucknow, India (26°50′N, 80°54′E) and reared in the laboratory. Adults were paired in plastic Petri dishes (9.0 × 2.0 cm) and fed on an *ad libitum* supply of pea aphid, *Acrythosiphon pisum* (Harris) from a *Vicia faba* Linnaeus host plant grown in a polyhouse (maintained at 22±3°C; 65±5% R.H. and 14L: 10D photoperiod). These Petri dishes were kept in BOD incubators (Yorco Super Deluxe, YSI-440, New Delhi, India) at 27±2°C; 65±5% R.H.; 14L:10 D and inspected twice daily (1000 and 1500h) for oviposition. The eggs laid were separated and kept in BOD incubator (abiotic conditions same as above). Each neonate larva was reared individually in Petri dishes (size as above) until adult emergence. Wild adult beetles were added to the laboratory stock every five generations to prevent inbreeding. These beetles, comprising unrelated individuals, were used for the experiment.

### a) Maintenance of inbred lines

Adults from both morphs were paired in Petri dishes, and their offspring were reared to maturity. Subsequently, these offspring were mated with their full siblings for five consecutive generations (Saxena et al., 2016). This resulted in an inbred line comprising individuals from both morph types that were siblings to each other. Related individuals from this line were used for the experimentation to study the effect of relatedness and to access the paternity since parental morph influences offspring phenotype enabling offspring paternity to be assigned (Chaudhary et al. 2017) (Plate 3).

### b) Double mating experiment

To investigate how relatedness affects the paternity share of the second male, we conducted various double mating experiments using typical female beetles (Plate 4). This approach was chosen based on prior research by Chaudhary et al. (2016), which eliminated other potential factors such as morph type influencing second male paternity share. In the first experiment, typical females were sequentially mated with an unrelated typical male followed immediately by an unrelated intermediate male after the first mating naturally concluded. In the second experiment, females were first mated with a related typical male and subsequently with a related intermediate male. The third experiment involved mating a typical female with an unrelated typical male followed by a related intermediate male. Lastly, in the fourth experiment, females were initially mated with a related intermediate male and then with an unrelated typical male. Each experiment was replicated 15 times to ensure robustness and reliability of the findings.

### c) Mating and reproductive attributes

Time of commencement of mating (from introduction in Petri dish to establishment of genital contact) and copulation duration was recorded for all the mating. After the second mating, females were isolated in Petri dishes (size as above) with *ad libitum* prey. Eggs were counted daily and their viability (number of larvae hatched from the eggs) recorded. Fecundity (average of eggs lay for seven days) and percent egg viability (the percentage of individuals hatched from the eggs laid) were recorded for each individual female. Each hatched larva was housed individually with prey until adult emergence. The percent of offspring sired by the related and unrelated males was recorded for seven consecutive days, and the overall paternity share was compared based on parental morph types. Parental morphs influence offspring traits and help determine paternity (Chaudhary et al., 2017). Pairs with the same morph produced offspring of that morph, while pairs with different morphs, like typical and melanic, produced offspring that were lighter than the melanic parent but darker than the typical.

### Statistical Analysis

All the data were analysed with IBM SPSS 20. The data were examined for normality and homogeneity of variances with the Shapiro-Wilk test and Levene’s test, respectively. A generalised linear model (GLM) was used to analyse the effect of status of second male (relatedness) on mating (time of commencement of mating and copulation duration) and reproductive parameters (fecundity and percent egg viability) using gamma distribution with mating and reproductive parameters as the dependent variables and effect of status of second male as predictors using different GLM models. Percent of paternity share was subjected to the chi square goodness of fit test by using MINITAB 20 software.

## Results

### Effect of mating order and male status (related or unrelated) on the mating parameters

Time of commencement of mating was not significantly influenced by mating order (χ^2^=1.56; df=1; P>0.05) and significantly influenced by the male status (χ^2^=17.10; df=1; P<0.05). The interactions between the two factors were also significant (χ^2^=15.66; df=1; P<0.05). Females took more time to commence second mating in the double mating treatments where the second males were related (Figure 1a). Copulation duration was also significantly influenced by mating order (χ^2^=34.72; df=1; P<0.05). The effect of the male status on copulation duration was insignificant (χ^2^=3.65; df=1; P>0.05). The interactions between the two factors were also insignificant (χ^2^=3.15; df=1; P>0.05). Females mated for shorter durations during second mating with related males as compared to the unrelated second male (Figure 1b).

**Figure 1:**
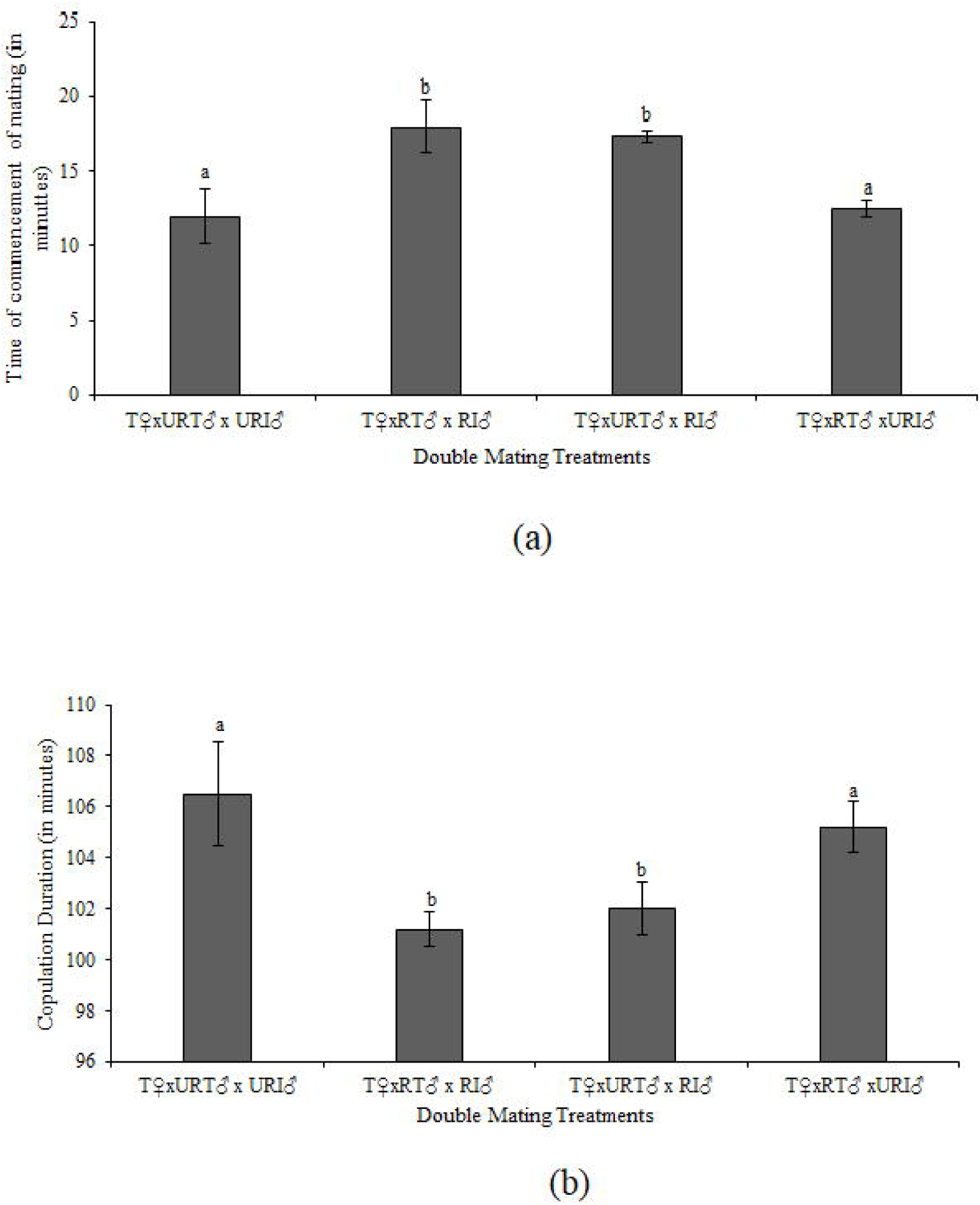
a) Time of commencement of mating and b) copulation duration of *Cheilomenes sexmaculata* when subjected to double mating treatments.

### Effect of status of first and second male (related or unrelated) on the reproductive parameters

The status of male have significant impact on the reproductive parameters. Fecundity was significantly influenced by the status of the first male (χ^2^=232.90; df=1; P<0.05). The status of second male was also significant (χ^2^=79.99; df=1; P<0.05). The interactions between the two was also significant (χ^2^=37.48; df=1; P<0.05). Females were more fecund when the second male was unrelated to the females (Figure 2a). Percent egg viability was insignificantly influenced by the status of the first male (χ^2^=0.00; df=1; P>0.05) and significantly influenced by the status of the second male (χ^2^=14.55; df=1;P<0.05). The interactions between the two were also insignificant (χ^2^=0.00; df=1; P>0.05). It was higher when unrelated second male was mated with females (Figure 2b).

**Figure 2:**
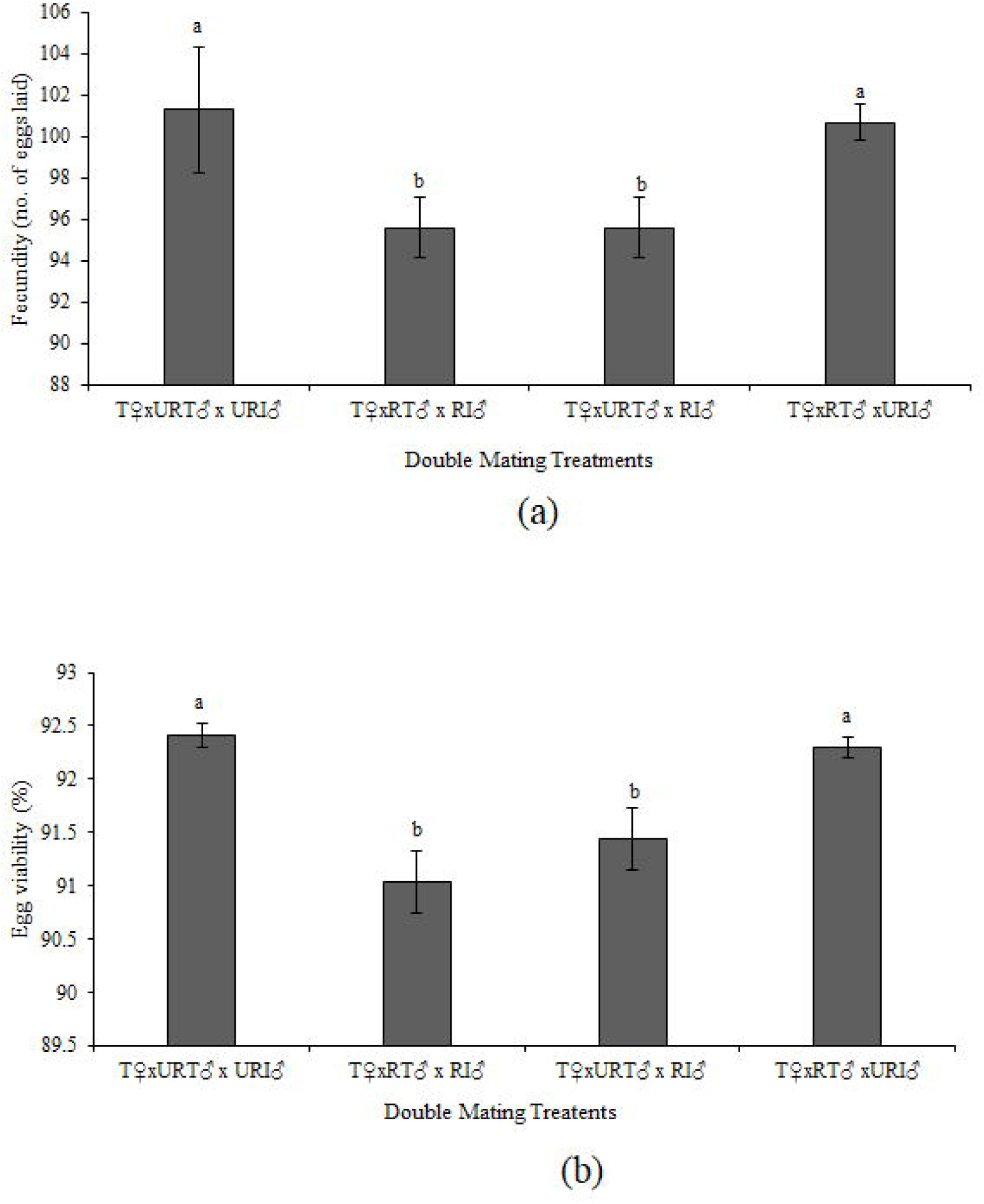
a) Reproductive parameters (a) fecundity (b) percent egg viability of *Cheilomenes sexmaculata* in double mating treatment with unrelated intermediate (URI) and related intermediate male (RI).

Percent paternity success of the second male was found lowest in the double mating treatment where the first male was unrelated and second was related to the female (Figure 3).

**Figure 3:**
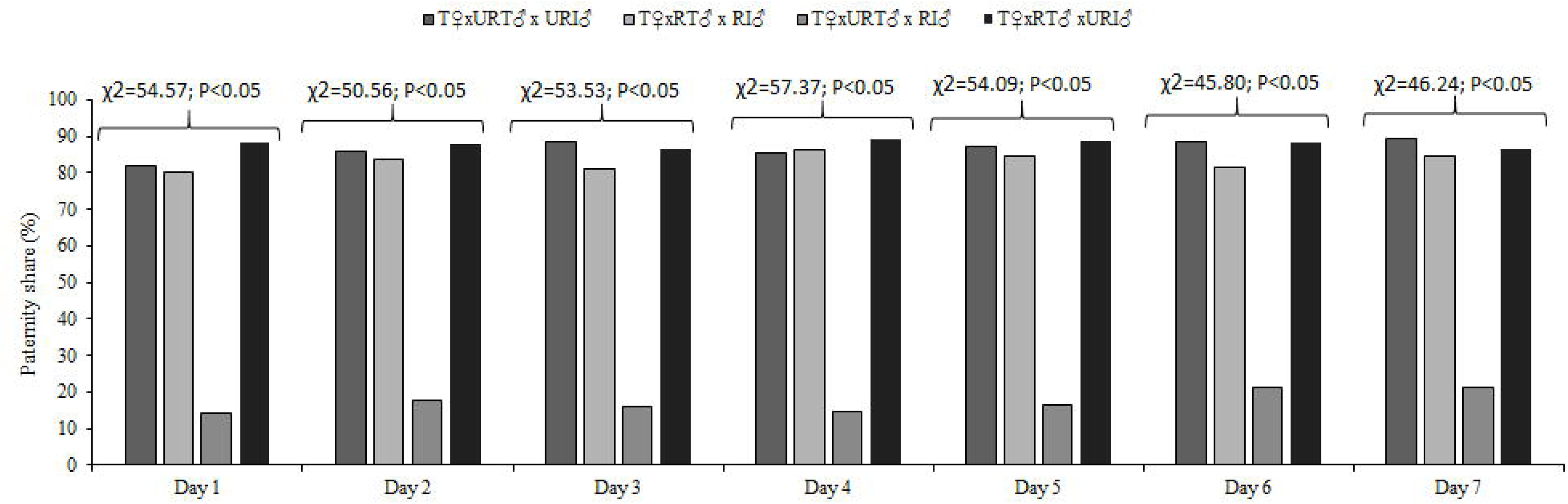
Percent paternity share of second males when females of *Cheilomenes sexmaculata* were subjected to different double mating treatments.

**Figure 4:**
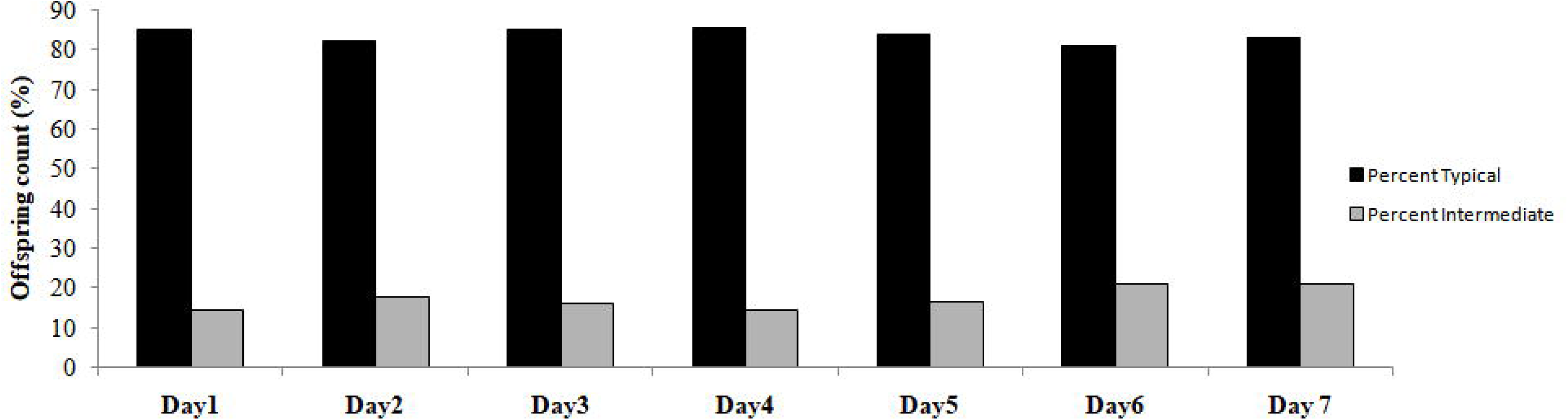
Effect of relatedness on morph of offspring of ***C. sexmaculata*** under double mating treatment with unrelated typical and related intermediate male (RI).

**Figure 5:**
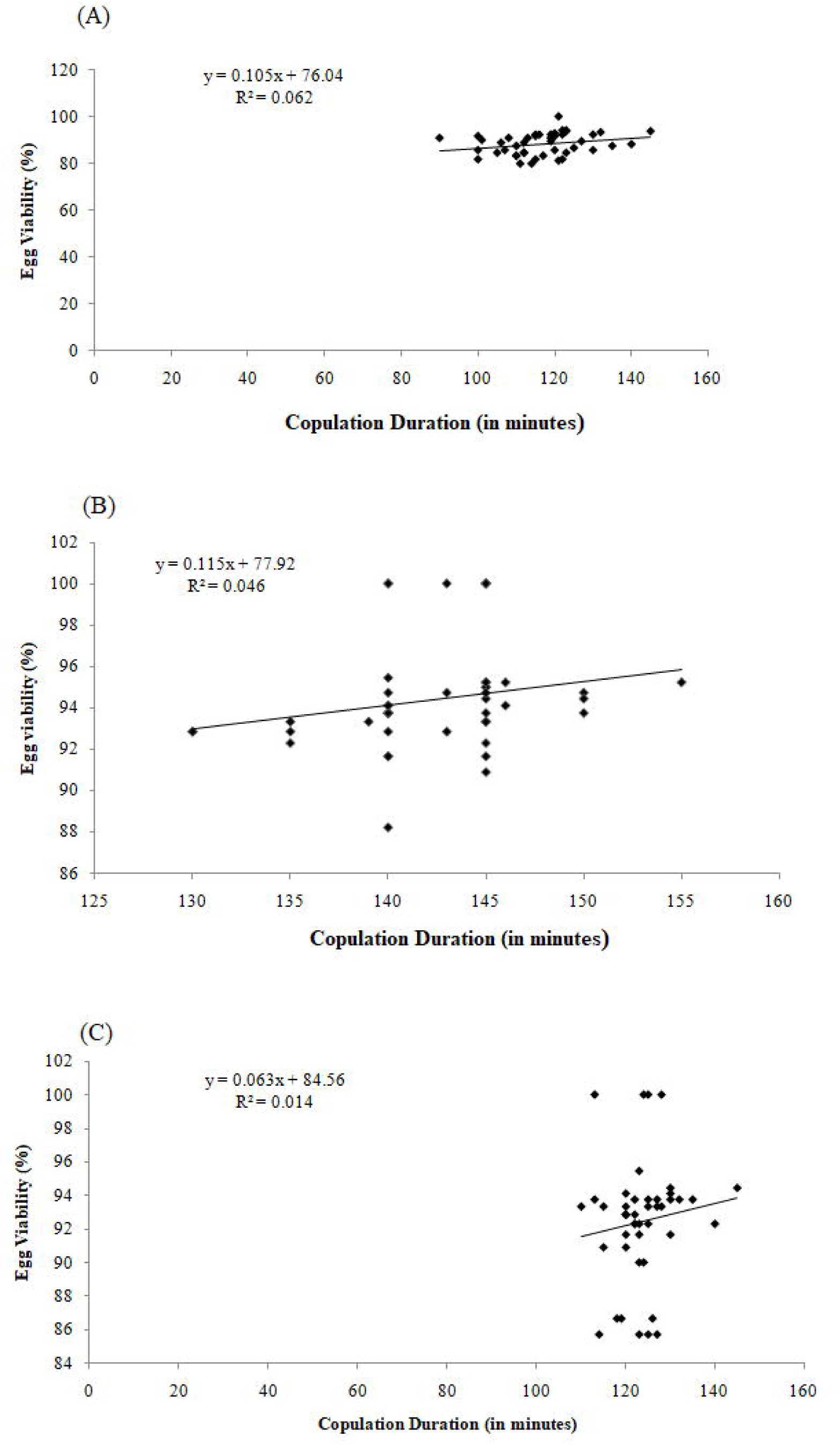
Relationship between copulation duration and percent egg viability in different sex ratio treatments when females mated with males reared on (A) Scarce Diet. (B) Abundant Diet. (C) Alternate Det

## Discussion

The present study revealed that the last male sperm precedence in *C. sexmaculata* is rather dynamic and breaks down when the last male to mate in related. In polygamous insects like ladybirds, sperms from different males compete for fertilization in female reproductive tract post multiple mating. This leads to an unequal sharing of paternity that favours the most recent male. This is termed as last male sperm precedence (LMSP). Previous studies have shown that in *C. sexmaculata*, last male to mate gains maximum paternity share, siring up to 72% of the offspring (Chaudhary et al. 2016). However, our study revealed that the last male sperm precedence in not static and is modulated with the quality of last male as relatedness in our case. When the second male was related (full sibling) to the female, the paternity tilted significantly in favour of the first unrelated male. Thus, in presence of sperms from the related male, the female may have preferred unrelated sperms over related.

Last male sperm precedence is relatively common in polyandrous insects (Simmons and Siva-Jothy 1993). There are commonly two mechanisms proposed to account for LMSP: (1) sperm displacement and removal; (2) stratification of sperm (Birkhead 2010). In damselflies, *Calopteryx maculate*, penises of males are covered with backward facing spines. This modification of penis aids in the removal of first male’s sperm from female bursa copulatrix and spermathecal tubes, ensuring maximum paternity of the last male (Waage, 1979). Sperm ejaculation via spermatophore ejection by female have been reported in ladybird beetle *Harmonia axyridis* (Obata and Hidaka, 1987). However, this was not observed in our case. According to sperm stratification, the last sperm to enter the sperm storage organ are the first to leave therefore yielding to the high paternity share P2 value (Simmons, 2001b). However, maximum paternity share of the first unrelated male when second male was related rules out the possibility of both, sperm displacement and sperm stratification. Higher paternity share of the unrelated male irrespective of its mating order (first or last to mate), indicates active role of the female over the process of sperm storage and suggests role of cryptic female choice. Possibility of related males ejaculating less sperms owing to reduced paternity share could not be denied and more detailed study is needed at molecular and physiological level to come to a firm conclusion.

Relatedness can modulate insect mating behaviour. For example, in the parasitoid wasp *Venturia canescens*, females avoid mating with siblings, to avoid the risk of producing sterile or inviable offspring (Metzger et al., 2010). Similarly, in *C. sexmaculata*, the increased time of commencement of mating and reduced copulation duration when mating with the related males, points to a pre-copulatory mechanism to recognize the genetically related (siblings) mate and avoid mating with them. The commencement of mating is the female governed phenomena while copulation duration is a male governed phenomena (Bind, 2007) and therefore it can be inferred that these beetles probably try to avoid mating with full siblings by either delaying onset of copulation or doing copulation for shorter duration of time (Saxena et al., 2016). Our findings suggests that other than reducing second male paternity success, relatedness of second male also affected time of commencement of mating which indicates the female acceptance of mating.

The ability of mate recognition is particularly common in insects (Hughes et al., 1999). Cuticular hydrocarbons (CHCs) are the most frequent form of chemical cues for recognition of potential mates (Lewis, 1984; Roitberg and Isman, 1992; Morgan, 2004; Tuni et al., 2013; Pattanayak et al., 2014; Gallot et al., 2020; Weiss and Schneider, 2021). For instance, crickets assess the relatedness of mates via CHCs (Tuni et al., 2013). Mate selection can be made at two levels (pre-and post-copulation in the form of sperm competition and cryptic female choice) therefore, the female did not show complete abstinence from mating with the related male, in order to avoid the cost of harassment by males (Singh, 2020). In *D. melanogaster* if the female fails to avoid mating with related males at precopulatory level then it uses her cryptic strategy to select the compatible male at postcopulatory level (Mack et al., 2002).

Prolong copulation duration is considered a crucial factor to ensure maximum paternity success (Chaudhary et al., 2015; Magris et al., 2020). In our study, copulation duration was significantly reduced when mating occurred between relatives and may be a response of mate avoidance with siblings. There are a number of other reasons potentially explaining the discrimination between the competing males. The “compatible genes” hypothesis correlates offspring fitness to genetic dissimilarity between mating partners and hence avoids mating with the siblings (Neff and Pitcher, 2005). Since mating between the genetically related individuals is likely to increase homozygosity, the risk of inbreeding depression is also enhanced (Müller and Juškauskas, 2018; Müller and Müller, 2016; Valtonen et al., 2014). To prevent such inbreeding depression, selection should favour kin recognition (Breed, 2014) and sibling avoidance in many species (Lihoreau et al., 2007; Fischer et al., 2015; Duthie et al., 2019). However, to see if the possibility of the avoidance is male driven, studies on ejaculate count/size and seminal fluid protein are required.

Earlier investigations in many taxa have been suggested that the females avoid to sire their eggs from the sperms of heterospecific males (Birkhead 2000; Alipaz et al. 2001; Price et al. 2001) and genetically incompatible conspecific males (Palumbi 1994; Olsson et al. 1996; Zeh and Zeh 1996, 1997; Alipaz et al. 2001). Since post copulatory selection gives the female, fitness benefits by choosing “good” and compatible mate (Firman et al. 2017; Tuni et al. 2013), reduced paternity success of the related male in comparison to that of the unrelated male could be the result of post copulatory sperm selection. Therefore, the female tends to influence the relative success of the ejaculates post mating, and modulate paternity success based on quality of male (Saxena et al., 2017).

Though there are several factors which affects the paternity success in ladybirds, in the present study, relatedness is used as the potential predictors of paternity success (either as the related first male or second male) and modulates the phenomenon of LMSP. The modulation of paternity may have evolved as a counter adaptation to male traits that promote last male sperm precedence (Laturney et al. 2018). The study also compels us to reconsider the mechanism of sperm stratification and investigate the possibility of sperm mixing and cryptic female choice in *C. sexmaculata*. The compromised reproductive fitness, possibly hints to cryptic female choice against related males. Therefore, the opportunity to avoiding mating with related individuals, is widely suggested to be a driver of adaptive evolution of multiple mating and sperm precedence (Varian-Ramos and Webster 2012; Kingma et al. 2013; Lehtonen and Kokko 2015; Reid et al. 2015; Duthie et al., 2016). Furthermore, assessing CHCs that could possibly play a role in the identification of siblings, is suggested, to determine any such pre copulatory recognition mechanism in *C. sexmaculata.* The mating bias exhibited by the female against the related males, could indelibly evolve the reproductive strategies as a means of inbreeding avoidance.

## Acknowledgements

SS is grateful to Prof. D.J. Hosken, University of Exeter, Exeter, UK, for improving the language of the manuscript and providing the valuable suggestions. SS also acknowledges BSR Fellowship by University Grants Commission, New Delhi, India (F. No.25-1/2014-15 (BSR)/7-109/ 2007/BSR), dated 25 August 2015.

## Data availability statement

Data generated or analysed during the study are available from the corresponding author upon reasonable request.

## Conflict of Interest

all authors are in agreement with the content of the manuscript and have no conflict of interest.

## Plates

Plate 1: Different types of morphs of *Cheilomenes sexmaculata*: (a) Typical, (b) Pure melanic, (c) Intermediate, and (d) Offspring of heteromorphic pair

Plate 2: Offspring morph types produced by Homomorphic and Heteromorphic pairs in *Cheilomenes sexmaculata*.

Plate 3: Development of inbred lines and utilization for paternity studies based on morph type Plate 4: Double mating treatments and the maximum percentage of offspring sired.

---

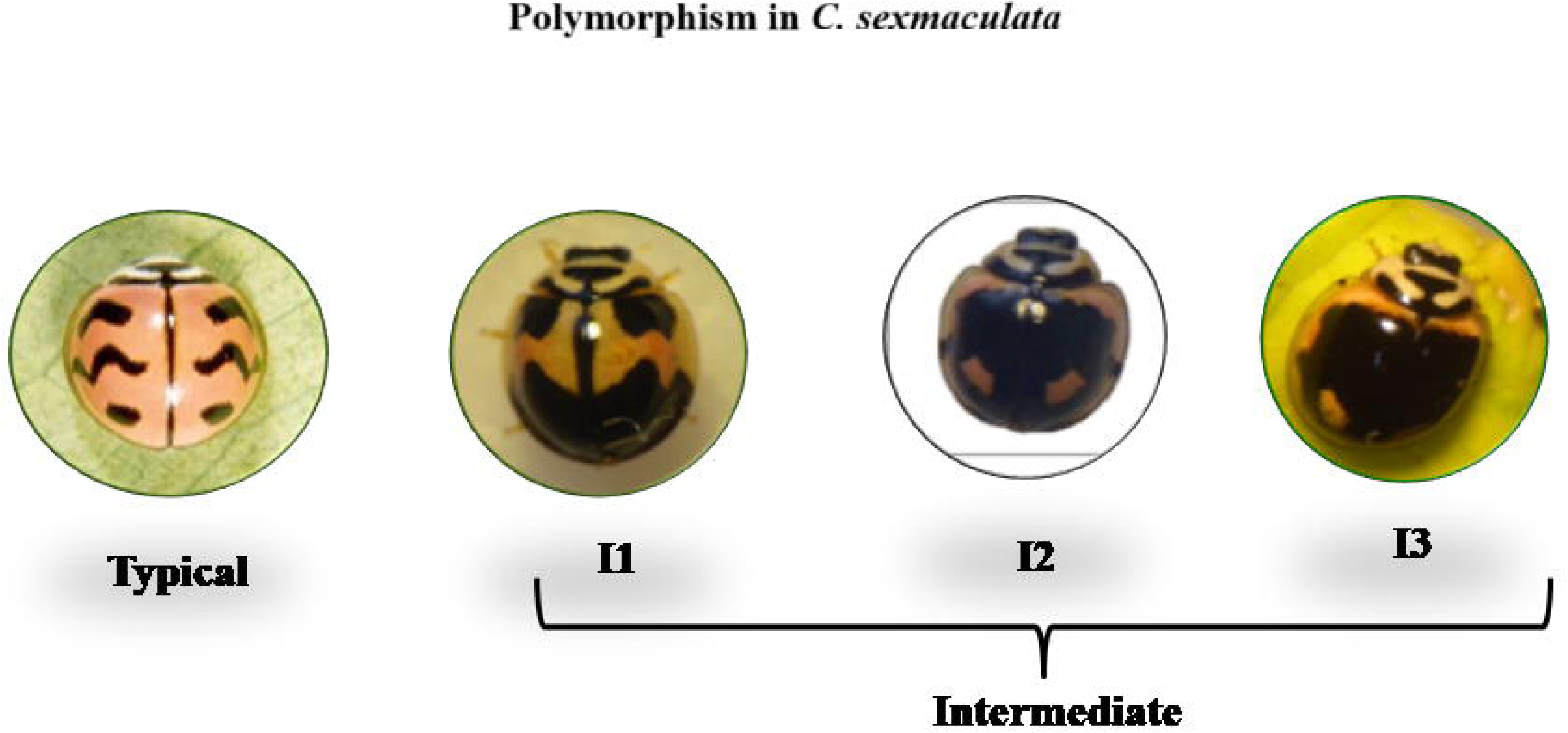

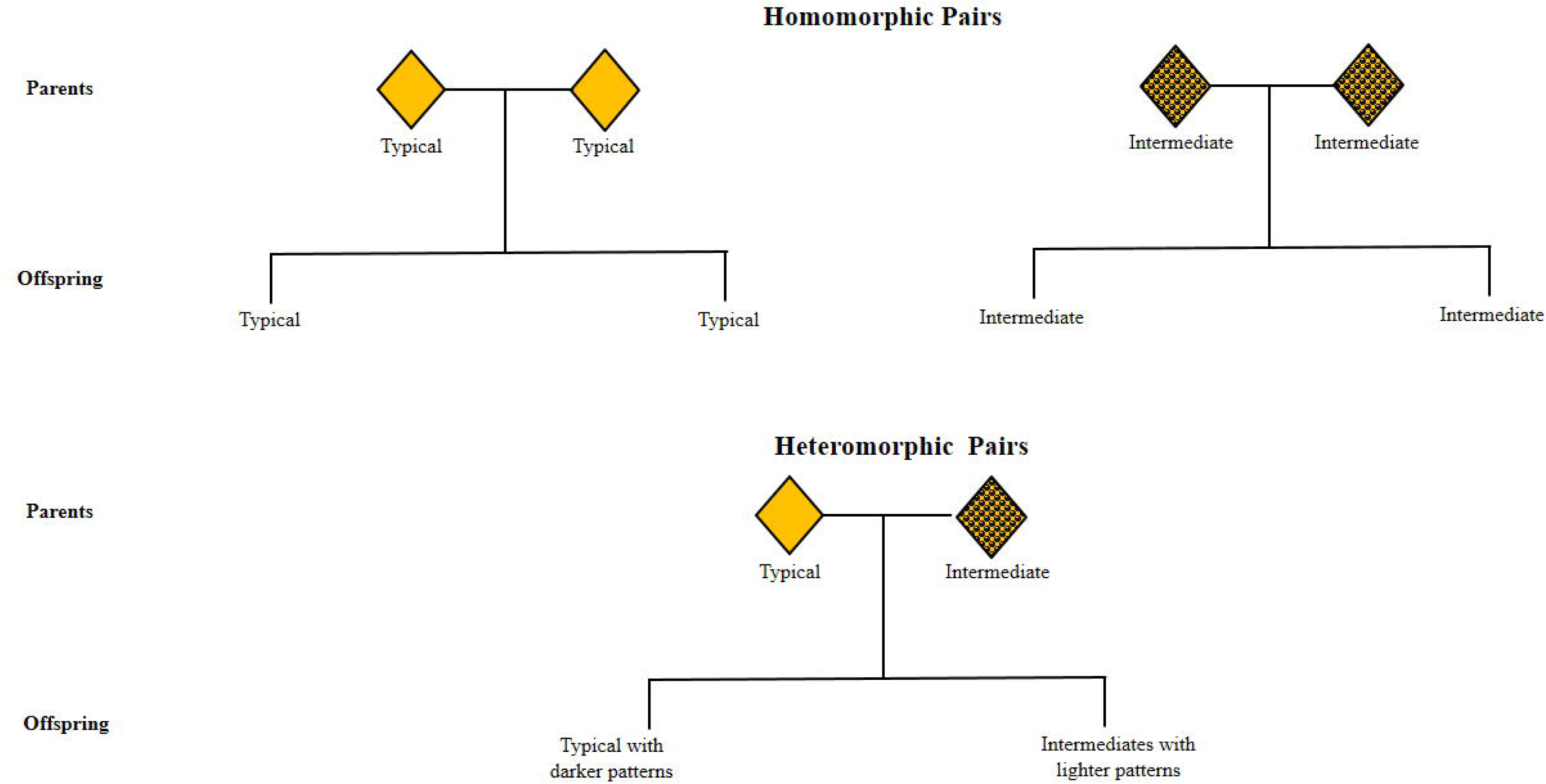

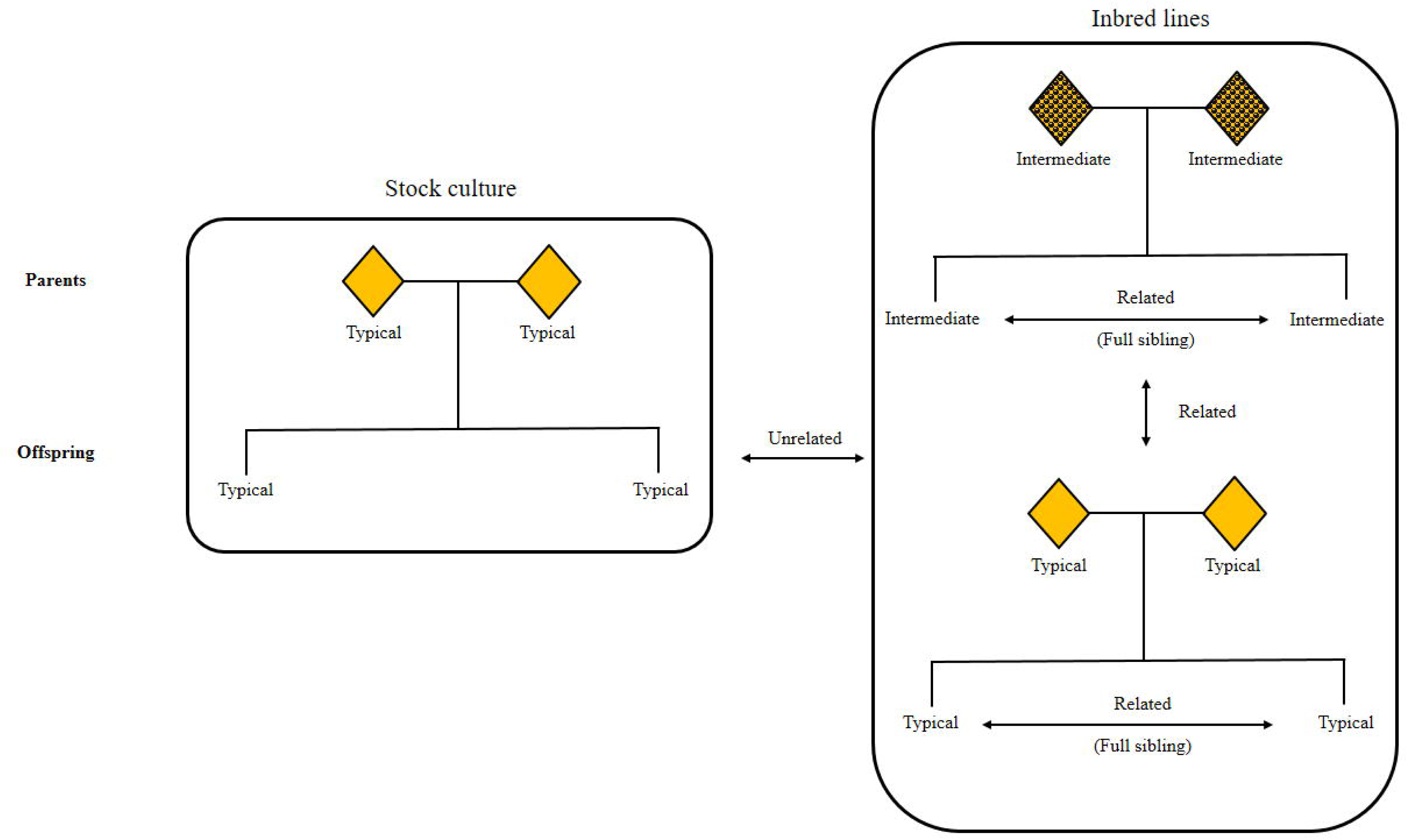

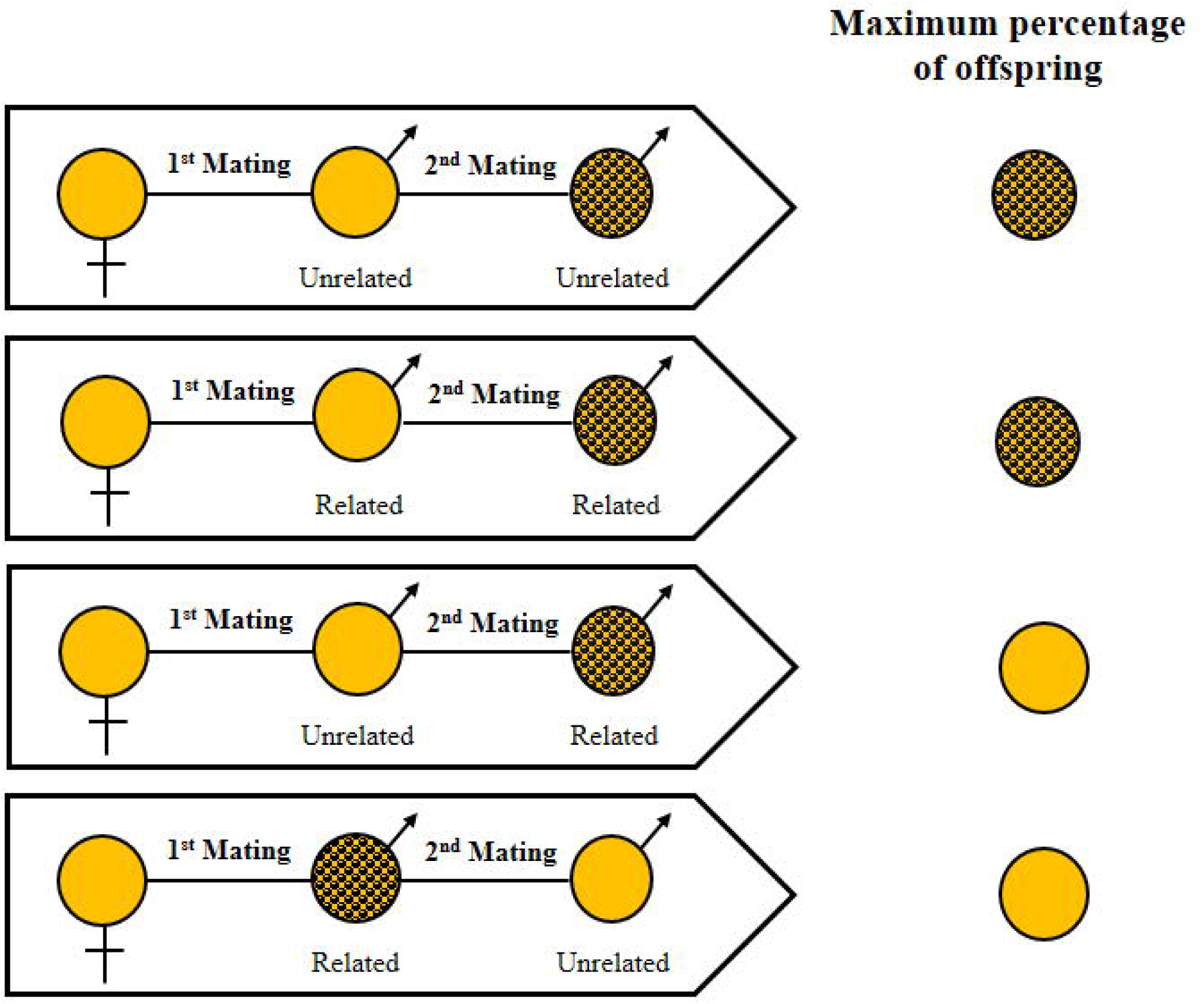

## Notes

### Competing Interest Statement

The authors have declared no competing interest.

